# ISCB RSG-Spain and highlights from the VIII Spanish Student Symposium in Bioinformatics and Computational Biology in 2021

**DOI:** 10.1101/2022.08.19.504447

**Authors:** Inés Rivero, Guillermo Jorge Gorines-Cordero, Luis A. Rubio-Rodríguez, Irene Soler-Sáez, Carla Perpiñá-Clérigues, Adrian García, Sara Monzón, Tamara Hernández-Beeftink

## Abstract

The Regional Student Group of Spain is recognized by the Student Council and the International Society for Computational Biology. The objective of these institutions is to connect and share useful information among all professionals and students working in the field of bioinformatics and computational biology. In this article, we intend to present and publicize RSG-Spain, the Spanish ISCB regional student group, by relating its recent history of a growing community of students and young professionals and how it helps their development in the field. RSG-Spain, since its creation, has been involved in the organization of events with the aim of gathering bioinformaticians in the country and promoting research and collaboration in the field. Here the VIII Student Symposium held online in 2021 is presented, analyzing the output of the event and showing the main challenges found in its organization.

## Background

The International Society of Computational Biology (ISCB) is a global society that gathers bioinformaticians around the world. The ISCB advocates for the promotion of grants, research, training, outreach, and inclusive community building in computational biology and its professions (https://www.iscb.org).

The ISCB has a Student Council (SC) in charge of promoting the development of the next generations of computational biologists (https://www.iscbsc.org). In 2006, the SC created the regional student group (RSG) program to endorse interaction among students pursuing research in the field of Bioinformatics and Computational Biology in different countries^1,2^. Currently, around 26 active RSGs comprise more than 1500 members worldwide. In particular, RSGs organize regional, national and even supranational activities and meetings to promote the professional development of students^3^. They also aim to serve as a bridge between the local industry and academia by providing career opportunities for the students.

### Regional Student Group of Spain

RSG-Spain was founded in 2010 by Lorena Pantano, a PhD student at the time, who discovered the existence of the ISCB SC and the RSG program in Boston. Almost single-handedly, she decided to start the Spanish RSG. Soon after, she managed to engage other PhD students, such as Salvador Capella, who is now our faculty advisor, to join this adventure. In the beginning, most members were from Barcelona, but since 2016 this network has expanded to other regions of Spain. In 2019, the network had enough members across the country to adopt the RSG-Spain local nodes system, starting with Barcelona (2016) the Canary Islands (since 2018), Madrid (since 2018) Granada (with BioInformaticsGRX association, https://bioinformaticsgrx.es/) (since 2018), and Valencia (since 2021), all of them being currently active. Some regions, like Cordoba or Esukadi, were also established in 2019 but are no longer active (**Figure 1**). RSG-Spain is always seeking students from other regions of Spain in order to make this network grow further and cover all the regions where our expert field is present.

**Figure 1.**
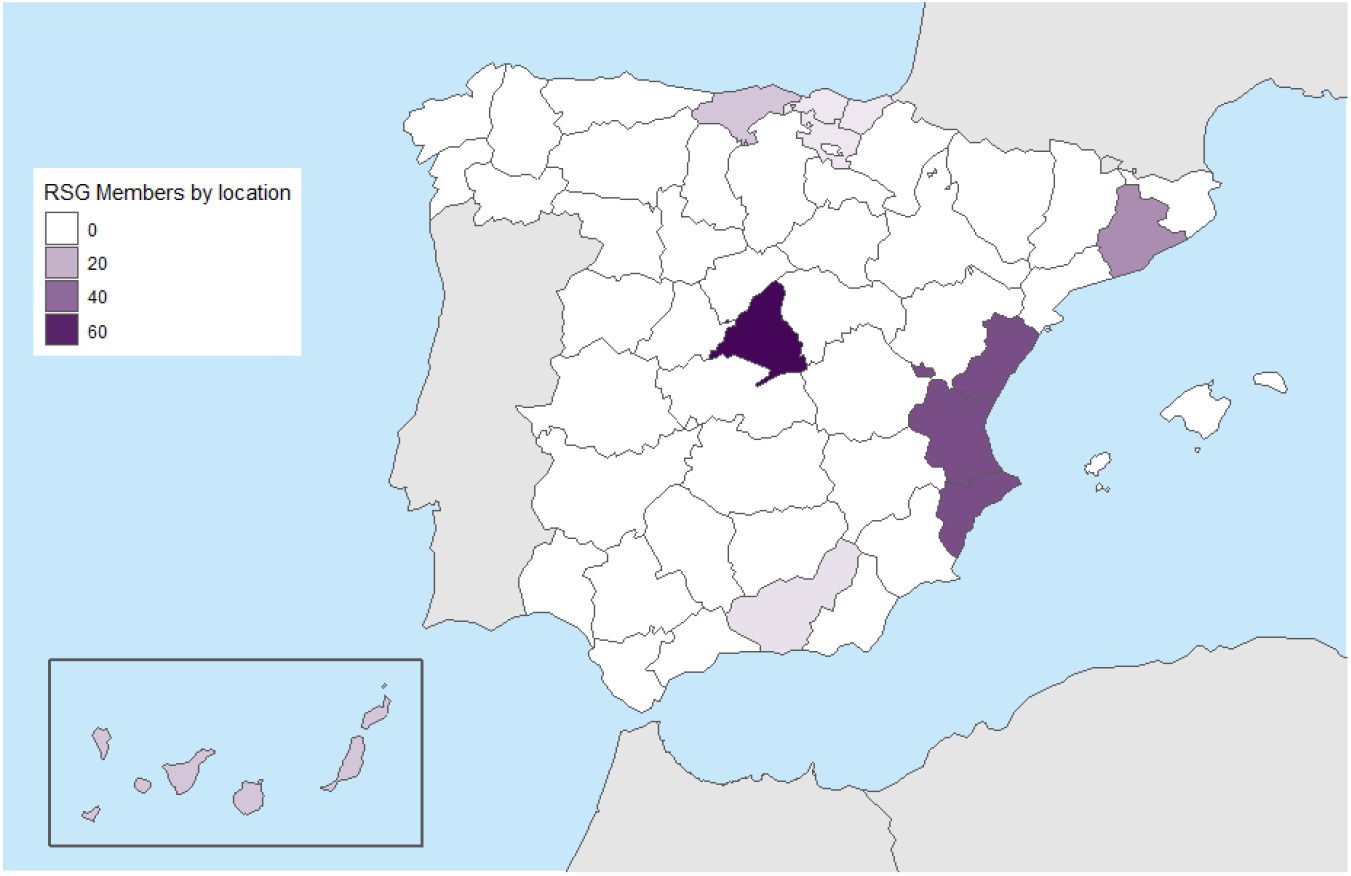
The population of each regional RSG inside of Spain as of June 2022.

Following an academic year, the RSG-Spain nodes individually and jointly organize monthly activities. Normally, for obvious reasons, the local nodes organize in-person events while leaving the national and international scope to the online platforms. Nevertheless, at least once a year, obviating the COVID-19 years, the core organizing team of RSG-Spain carries out an in-person symposium as a satellite meeting of the senior professional’s congresses that can be national or international. These are organized with an association routine of local and general meetings, on which projects and events are planned, and useful information is shared with our fellow members.

Our community is kept updated with a constant feed of bioinformatics job opportunities, divulgative talks, web blogs, scientific seminars, highlighted articles and tutorials by using social media and our mail distribution list. Furthermore, the group is always willing to organize workshops, social networking events and scientific sessions, trying to provide awards and recognition for the outstanding contributions presented at the student symposium. As a result, students share their experiences and concerns which help ease the development, personally and professionally, as computational biologists and bioinformaticians. As part of our outreach, different social media channels are maintained, including Twitter (@RSGSpain), Instagram (@rsg_spain), YouTube channel (ISCB RSG-Spain) and our own website (https://www.rsg-spain.iscbsc.org) to share all this not only with our members but also with the whole scientific community.

### Previous Symposia

Since 2010, we have organized several Student Symposiums. Starting in 2010 with the I Student Symposium, which took place as an initial session of the “*Jornadas de Bioinformática*” in Málaga, and was arranged together with Portugal and North-Africa Regional Groups. In 2012, the second edition of this event took place in Barcelona, also together with Portugal and North Africa. In 2014, the third edition symposium took place in Sevilla. And our subsequent editions were held in Valencia, Barcelona and Granada, in 2016, 2017, and 2018 respectively. In 2019, our VII Bioinformatics Student Symposium was co-organized by INB (https://inb-elixir.es)^4^ and Centro de Biotecnología y Genómica de Plantas (CBGP, https://www.cbgp.upm.es). It gathered more than 90 students in Bioinformatics from around Spain. The conference was held on 17th October 2019 in Madrid, as a satellite event of the *“II Jornadas de Bioinformática y Biología Computacional de la Comunidad de Madrid” (II BioinfoCAM)* organized by CBGP (UPM-INIA).

### VIII Student Symposium (2021)

On the 18th and 19th of October 2021, we held our VIII Bioinformatics Student Symposium. This time, due to the pandemic, the event was entirely held online. For this purpose, we made use of the Airmeet platform (https://www.airmeet.com), which was provided by the ISCB-SC. The main objective of this event was to generate and encourage debate and collaboration between our network, also promoting RSG-Spain, the SC and the ISCB.

On the first day, we held a Meet & Greet event, where we presented the RSG-Spain organization and the respective active nodes in 2021 (Barcelona, Granada, Madrid and the Canary Islands) to the attendees. We showed our main activities at national and local levels, as well as we promoted the creation of new nodes throughout the Spanish territory. Also, we carried out a survey of all the young researchers who attended the event on Bioinformatics topics and practices in Spain, which gave us a general perspective of the current panorama in our country. Moreover, this day we had a Round Table at which different bioinformatics students discussed the ongoing Spanish trends in bioinformatics and computational biology. We also had a high-quality informative event, such as the splendid talk given by Carla Padilla (current president of RSG-Argentina) about how bioinformatics, and especially students, are performing currently in her country. Subsequently, from this beautiful intervention grew the idea of holding a student congress on bioinformatics in Spanish with the objective to eliminate access barriers to communication and presentation of international scientific papers (https://seh2bioinfo.netlify.app/).

On the second day, we celebrated the so-called Scientific Meeting, with very outstanding talks, such as the masterful and inspiring presentations by PhD Arcadi Navarro and PhD Laura Furlong. In these talks, they exposed their points of view and the different ways in which they came to make their incredible research career hand in hand with bioinformatics. On this second symposium day, we also had the participation of Nataly Bulson (BIOINFO 4 WOMEN, https://bioinfo4women.bsc.es), who spoke to us about gender perspective in research.

A total of 38 scientific communications from levels ranging from final degree projects to thesis were presented, of which 10 were previously chosen as oral communications, and other 10 as lighting talks (**Figure 2**).

**Figure 2.**
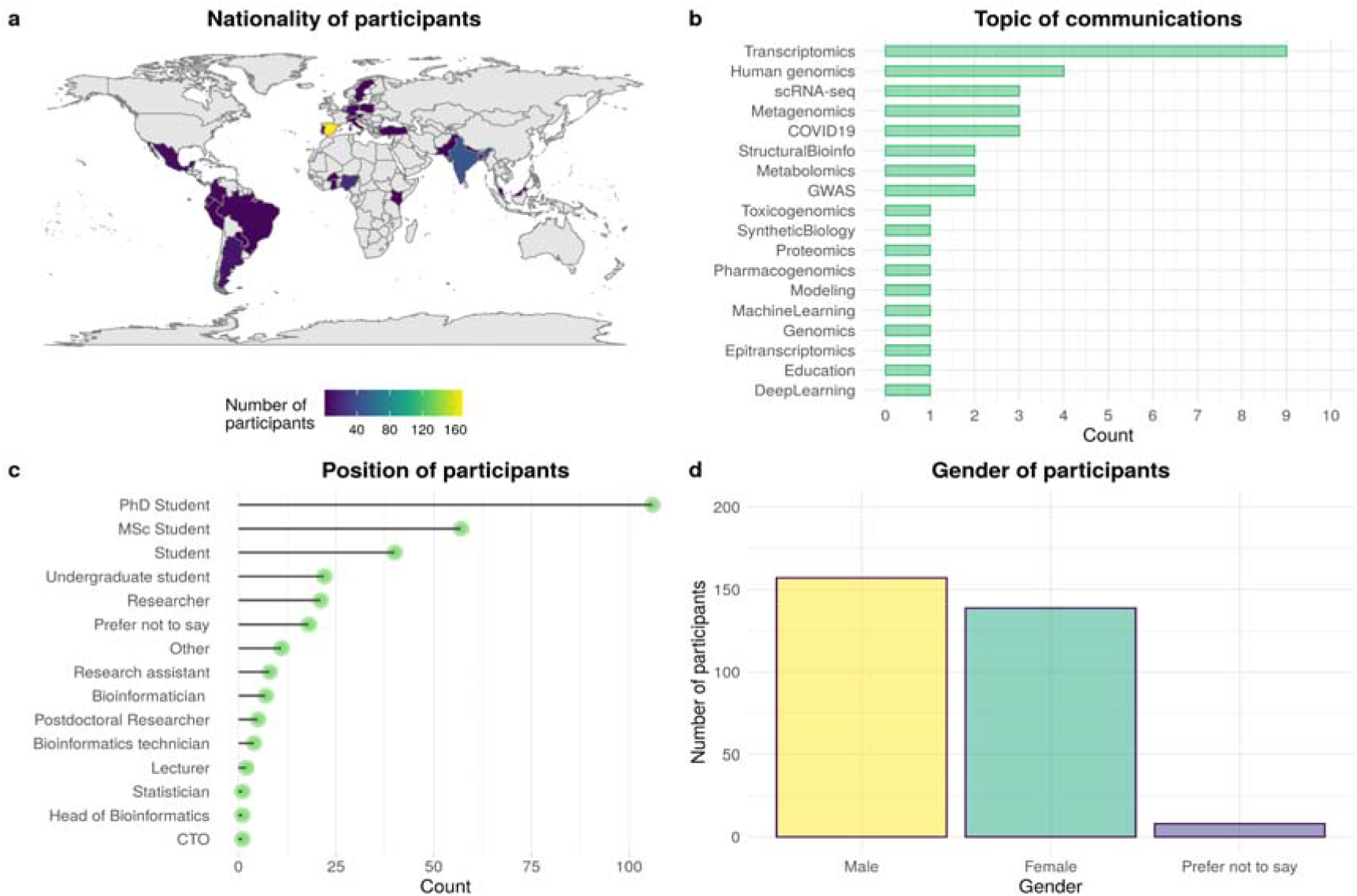
The VIII Bioinformatics Student Symposium in numbers. **a)** Map summarizing the country of origin of participants in the VIII Bioinformatics Student Symposium. Attendees from a total of 32 different countries registered for the event, being Spain the most predominant country of origin, with 169 of 304 attendees. **b)** Barplot summarizing the main field of the submitted communications. A total of 38 communications were submitted, of which 17 were presented as posters, 10 were presented as a poster with lighting talk and 10 were presented as a talk. **c)** Summary of the professional status or position of the attendees. From a total of 304 attendees, over one-third were PhD students, followed by Master’s and Undergraduate students. **d)** Barplot summarizing the gender of participants in the VIII Bioinformatics Student Symposium.

The VIII Spanish Bioinformatics Student Symposium proved to be a meeting point for more than 100 bioinformatics students and professionals from Spain and abroad, such as India, the USA, Malaysia and the Netherlands, among others (**Figure 2**). The level of scientific communication was remarkable, many exciting questions were asked, and many discussion topics were issued. These bouncy interactions showed that the field is more alive than ever in our country. For this Symposium, we had News In Three Lines (NITL, https://nitl.app/), Spanish National Bioinformatics Institute (INB, https://inb-elixir.es/) and the ISCB-SC as sponsors.

Furthermore, during these days, the participants and attendees were able to interact with the speakers and discuss their scientific work remotely from anywhere in the world. This demonstrates the usefulness of the online platforms for such events, and the great progress that moving to the online world has meant due to the pandemic situation.

### Summary of survey results during the Meet & Greet event

During the survey of our Symposium, all the young researchers who attended the event were able to discuss, summarize and obtain a general perspective of current bioinformatics in our country. This survey was mostly attended by students and professionals from different institutions throughout Spain, although there were also some participants from India, the United States, the Philippines, Bosnia and Herzegovina, Egypt, and Malaysia.

In general, there was an equal representation of males (46%) and females (50%), being 4% non-binary. Most participating students were between 22 and 25 years old (53%). The majority of the participants were PhD students (59%), followed by master’s students (35%), undergraduates (4%) and post-doctoral (2%), whose primary background came from Biology (Biotechnology, Biochemistry and more Bio-type) (86%), followed by Engineering (4%), Mathematics (2%), Physics (2%) and Informatics (2%). Likewise, we observed that most of the students began to study bioinformatics through regulated education (degrees) (60%), while the rest were self-taught (26%) and learnt by taking specific courses (14%). Linked to this, most participants prefer to learn bioinformatics through regulated education (degrees) (34%), in-person courses (24%), online courses (22%), web surfing (forums Stack Overflow, BioStars and Google) (18%) and to a lesser extent through online books (2%), where the participants also suggested favourite resource stories such as Google, BioStars, Stack Overflow or Github issues, and the 80% stated that they normally take additional bioinformatics courses/workshops. In addition, their main application areas were biomedicine (58%), microbiology (13%), software development (7%) and basic biology (7%), among others.

Interestingly, most of the participants did not believe that in order to work in a private company you need to have a doctorate (60%). The 25% of the participants declared to work in the field of bioinformatics and to have a monthly salary range between 1,000-1,300 euros, followed by 1,300-1,600 (23%) or no salary (23%), whose salary mainly came from public scholarship (31%), research project (29%) or public contract (26%).

Furthermore, the students reported using R (64%) and Python (19%) programming languages the most, but they preferred using Python (52%) instead of R (26%) or Perl (17%). Moreover, most of the participants preferred Linux and similar operating systems (78%), followed by Windows (17%) and to a lesser extent the Mac system (5%). While the text editor preferred by users is RStudio (22%) or Visual Studio Code (18%) and the rest used other editors like Sublime Text (11%), Jupyter Laptops (11%), Atom (11%) or Nano4 (9%).

During the survey, other curiosities were asked, such as the knowledge and use of GitHub or GitLab, how many screens they use to work or would like to have, as well as the use of a personal computer or laptop to do their job. As expected, most participants knew (93%) and used (69%) GitHub or GitLab. In addition, 79% of the students declared using their personal laptop or computer for work reasons with at least one (45%) or two screens (52%). In addition, 57% showed that their work or practice institution does not provide a personal laptop and computer, and 61% of the participants do not prefer to work and study with their own laptop, but fortunately, the majority of them have a proper workplace and set up to work with a computer (89%).

Another interesting topic covered in this survey was the great importance of mental health in students. In this section, the doctoral students stated that in general, they felt more or less hopeless and equally empowered (22%), but the majority specified that in their first year of doctorate they felt more empowered (27%) than hopeless. But as the years of doctoral studies progress, we observed that students begin to feel more hopeless. In fact, the students shared that they have suffered different mental health issues such as imposter syndrome in 84%, anxiety in 65%, social anxiety in 32% and depression in 32%. Finally, the participants considered that the most important factors for good mental health during the PhD are mostly to have work-life balance (76%), mentor guidance (57%), work environment (51%) and funding stability (41%). These results are quite interesting given the importance that mental health is currently gaining as it has been highlighted by several research studies in different countries around the world^5-6^. Linked to this, these observations have become more relevant and evident after the significant impact of the COVID-19 pandemic on the mental health of doctoral students^7-9^.

## Conclusions

After this symposium, many people joined our network, to the point that we have even recently accomplished one of our main objectives: to create a new local node in Spain (Valencia Node). The online interaction and participation have enriched our symposium and our RSG-Spain community. Thanks to this, we have more visibility and we are increasing our visits and views from most countries around the world. We have expanded our communication network and have developed valuable relationships that will certainly allow us to grow as professionals and engage in new projects. Despite this success, the main drawback is the inability to meet in person and hold networking events that enrich and create, among other aspects, closer friendships and interactions. Therefore, combining both worlds seems to be a good idea: including the social as well as the expansion part to become a richer community and make our network known.

## Acknowledgements

Thanks to all the team and students who are and have been part of RSG-Spain, and the VIII Student Symposium (2021) organizing committee.

## Author contributions

IR, GGG, LRR and THB prepared the manuscript. IR and GGG prepared figures. IR, GGG, LRR, ISS, CPC, AG and SM reviewed the manuscript. THB reviewed, finalized and communicated the manuscript. All authors collaborated to organize the event using the International Society for Computational Biology’s Student Council as a platform.

